# Heterogeneity of Transcription Factor binding specificity models within and across cell lines

**DOI:** 10.1101/028787

**Authors:** Mahfuza Sharmin, Héctor Corrada Bravo, Sridhar Hannenhalli

**Author notes:** **Corresponding author**: Sridhar Hannenhalli, 3104G Biomolecular Sciences Building (#296), University of Maryland, College Park, MD 20742, USA (v) (f).

## Abstract

Complex gene expression patterns are mediated by binding of transcription factors (TF) to specific genomic loci. The *in vivo* occupancy of a TF is, in large part, determined by the TF’s DNA binding interaction partners, motivating genomic context based models of TF occupancy. However, the approaches thus far have assumed a uniform binding model to explain genome wide bound sites for a TF in a cell-type and as such heterogeneity of TF occupancy models, and the extent to which binding rules underlying a TF’s occupancy are shared across cell types, has not been investigated. Here, we develop an ensemble based approach (*TRISECT*) to identify heterogeneous binding rules of cell-type specific TF occupancy and analyze the inter-cell-type sharing of such rules. Comprehensive analysis of 23 TFs, each with ChIP-Seq data in 4-12 cell-types, shows that by explicitly capturing the heterogeneity of binding rules, *TRISECT* accurately identifies *in vivo* TF occupancy (93%) substantially improving upon previous methods. Importantly, many of the binding rules derived from individual cell-types are shared across cell-types and reveal distinct yet functionally coherent putative target genes in different cell-types. Closer inspection of the predicted cell-type-specific interaction partners provides insights into context-specific functional landscape of a TF. Together, our novel ensemble-based approach reveals, for the first time, a widespread heterogeneity of binding rules, comprising interaction partners within a cell-type, many of which nevertheless transcend cell-types. Notably, the putative targets of shared binding rules in different cell-types, while distinct, exhibit significant functional coherence.

## Introduction

Transcriptional regulation is critically mediated by the binding of transcription factors (TF) to specific DNA elements in the genome (JACOB & MONOD 1961; Busby & Ebright 1994). While the *in vitro* binding specificity of many human TFs has been determined, it is well recognized that the *in vitro* binding specificity of a TF does not explain its condition-specific *in vivo* binding specificity (Zinzen et al. 2009; Yáñez-Cuna et al. 2012). This recognition has spurred investigations of additional determinants of *in vivo* binding, such as heterogeneity of TF’s binding motif (Hannenhalli & Levy 2002), homotypic clusters of binding sites (Dror et al. 2015), cooperative binding of the TF with its partners (Wang et al. 2006), condition-specific chromatin context (Heintzman et al. 2009), local DNA properties (Dror et al. 2015), epigenomic context (Gheldof et al. 2010) etc. While overall, both local genomic and epigenomic features have been deemed important in determining in *in vivo* occupancy of a TF, recent reports suggest that *in vivo* binding of a TF can be accurately predicted based only on the genomic signatures near the binding site (BS) without relying on the epigenomic context (Arvey et al. 2012; Dror et al. 2015); this is consistent with very recent reports showing that the epigenome itself is encoded by the genomic context (Whitaker et al. 2015; Benveniste et al. 2014). Taken together, these results strongly suggest that proximal genomic elements are the primary driver of *in vivo* TF binding. Prior sequence-based models of *in vivo* TF binding have shown that, somewhat counter-intuitively, the genomic context of a BS, which is the property of the genome, effectively encodes the condition-specific *in vivo* binding specificity (Arvey et al. 2012). This can be explained by the substantial plasticity of a TF’s interaction with other TFs’ and the modular nature of TF binding cooperatively with other TFs (Frietze & Farnham 2011), such that availability of specific combination of interacting TFs can guide *in vivo* binding to specific loci where the BS of the interacting TF are present in close proximity to each other, along with the availability of corresponding TFs (Hannenhalli & Levy 2002).

Previous sequence-based modeling of *in vivo* TF binding was done in a cell type-specific fashion. These cell type-specific models exhibit substantial *inter-*cell type heterogeneity, as expected, given variation in the availability of the potentially interacting TFs. However, these previous approaches build a single model for a cell type, thus implicitly assuming a homogeneous cell type-specific model, and as such have not investigated *intra*-cell-type model heterogeneity. Such heterogeneity of TF binding ‘rule’ across the genome can be expected for the same reason as for the inter-cell type heterogeneity. Moreover, in many instances, a binding specificity model trained in one cell type can predict a subset of *in vivo* binding in a different cell type (Arvey et al. 2012), suggesting that models of binding, or parts thereof, may be shared across cell types. Overall, the heterogeneity of sequence-based models of cell type-specific *in vivo* TF binding, and the extent to which a subset of binding rules (*sub-models*) are shared across cell types, is not known, motivating the present study.

To this end, we have developed an ensemble model based approach (***TRISECT***) to reveal both cell-specific and cell-independent rules for *in vivo* TF binding. We applied *TRISECT* to 23 TFs, each with genome-wide *in vivo* binding data in 4 - 12 cell types (a total of 135 TF-cell type combinations). For each TF, for each cell type, we built ensemble models of *in vivo* TF binding (***EMT***), then decomposed each *EMT* model into sub-models and clustered the pooled set of sub-models across all cell types using feature selection. Our comprehensive analyses strongly suggest that the cell type-specific binding rule for a TF consists of multiple sub-models, supported by our result showing that *EMT* captures the binding specificity better than previous non-ensemble models (Arvey et al. 2012). Moreover, for many TFs, the sub-models are shared across cell-types, and interestingly, we found that the putative target genes for similar sub-models across cell types exhibit a high degree of expression and functional coherence, suggesting that the *in vivo* binding rules are related to function of the gene targets, much more so than the cell type they are derived from.

In further probing the superior performance of *EMT*, we demonstrate that while a model based only on the known motifs of the reference TF, *i.e.* without incorporating additional potential TF interaction partners (*NonInteraction* model), can predict *in vivo* binding with ~78% accuracy, when motifs for other TFs are used in the model (*Interaction model*), the prediction accuracy is substantially increased to over 90%. Moreover, we found that the improvement in prediction accuracy by the *Interaction* model strongly correlates with the increase in the number of interaction partners, i.e., with model complexity, suggesting that the *Interaction* model effectively captures the heterogeneity of the binding rule. We identified and validated, based on literature, the potential interaction partners (we will refer to these as co-factors) that mediate context-specific binding and function of a TF. Finally, we show that certain TFs with multiple distinct binding motifs prefer binding to different motifs in different cell types, which may in part be associated with their inter-cell type variability of co-factors (Slattery et al. 2011).

In sum, our analysis reveals distinct sub-models of *in vivo* TF binding within a cell type that are nevertheless shared across cell types, and the shared sub-models across cell types target distinct yet co-functional genes in different cell types. A refined understanding of the genomic context of *in vivo* binding specificity can facilitate future investigations of transcriptional regulation and understanding of its genetic determinants.

## Results

### TRISECT – *Ensemble model of TF binding and the Clustering of sub-models across cell types*

#### Overview

The full analysis pipeline, *TRISECT*, is illustrated by Fig 1A. As the first step, we developed an ensemble model (*EMT*) to discriminate a TF’s *in vivo* bound genomic loci from the background, balancing model complexity (number of sub-models in the ensemble) against the cross-validation classification accuracy. Given a set of genome-wide loci bound by a specific TF, we first construct a foreground set of sequences (100 bps) centered at the ChIP-Seq peak. As a stringent background control, as done previously (Arvey et al. 2012), we use 100 bps regions ~200 bps away from the peak location (M&M). We considered a variety of feature sets for discrimination (see below). The *EMT* model was trained using Adaboost method where each sub-model is a decision tree built from a bootstrap sample (Friedman et al. 2000; Friedman 2002; Freidman 2008). Next, for a TF, given EMT models for all cell types, we represented each cell type-specific sub-model as a point in a *d*-dimensional space corresponding to *d* selected features (M&M). We clustered the data points, representing all sub-models in all cell types considered for a TF, using XY-fused network (*XYF*) (Melssen et al. 2006) such that sub-models within a cluster represent similar binding rules, either within a cell-type or across cell types.

**Figure 1:**
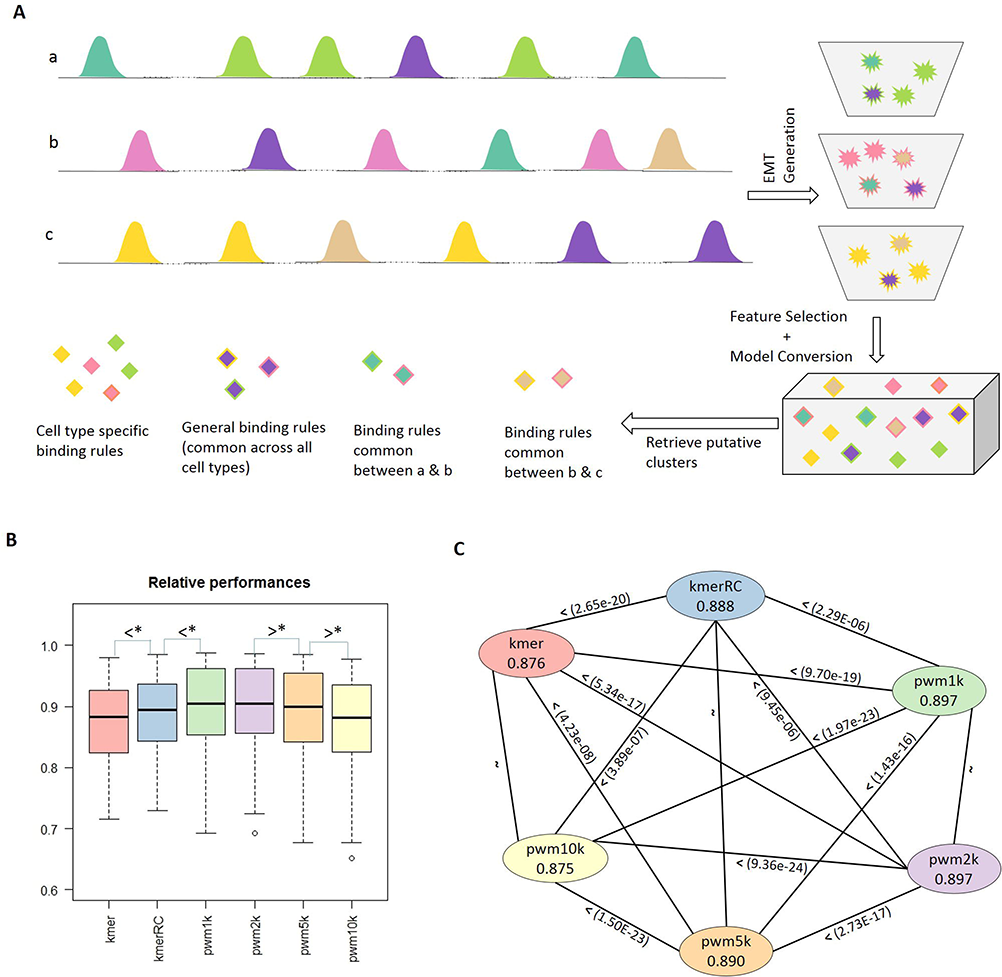
(A). Schematic of *TRISECT* pipeline. Different colors represent different binding rules or sub-models. Rows (a, b, c) represent cell types. Green, pink and yellow colors indicate cell type-specific sub-models. Each EMT is represented by a bucket of sub-models (top right). Star denotes sub-models and diamond denotes the corresponding data point after transformation into reduced feature space. The sub-models across all cell types are clustered. Cyan is common between cell types *a* and *b*, light-brown is common between cell types *b* and *c*, and purple is common across all three cell types. (B). Accuracy (ROCAUC) distribution for 6 choices of feature sets for EMT. (C). Comparison of accuracy between all pairs of 6 feature-set choices. Nodes are labeled with feature type and mean accuracy. Directional edges are labeled with Wilcoxon p-value.

#### EMT Feature sets

We considered three types of feature sets for a 100 bps sequence – (i) Kmer: frequency of occurrence for all 4096 6-mers, (ii) KmerRC: frequency of occurrence for all 2080 6-mers where a k-mer and its reverse complement were unified, and (iii) aggregate binding scores for 981 vertebrate TF motifs from TRANSFAC database (we used four stringencies for motif match) (M&M); we refer to these as *pwm* models. We applied *TRISECT* to 23 TFs, each with ChIP-Seq data in 4 to 12 cell types (a total of 135 TF-cell pair *EMTs*), listed in Supplementary Table 1. ATF was included in this study if (i) TF has narrow-peak data for at least 4 cell lines with at least 4k sites in each cell line, and (ii) TF has established PWM in TRANSFAC 2011 database. The performance assessment of *EMTs* was conducted based on 25% held-out dataset. The overall performance is summarized in Fig 1B and details are provided in Supplementary Table 2.

#### EMT performance

Fig 1B shows the overall accuracy distribution (over 135 TF-cell type pairs) for the 6 types of models, where the accuracy is quantified using ROCAUC on the test set. We compared the performances, using Wilcoxon test, among 6 sets of *EMTs* (kmer, kmerRC, and PWM at 4 stringencies) containing 135 TF-cell type pairs in each set (Fig 1C). We found that kmerRC significantly outperforms kmer model (Wilcoxon p-value 2.65E-20), consistent with the fact that TF binding occurs on double-stranded DNA and as such does not have directionality (except in relation with other interacting TFs) and therefore unifying each kmer with its reverse complement provides a better abstraction of biological determinants of TF binding. Following this line of reasoning, PWMs provide an even better abstraction of DNA binding specificity and as expected, the PWM-based models outperform kmer-based models (p-value, 2.29E-06 comparing kmerRC and pwm1k). Based on relative performances we selected pwm1k-based EMT for feature selection and clustering of sub-models and all subsequent analyses.

#### Comparison with previous model

Next, we compared *EMT* model (using kmerRC and pwm1k) with previously published model based on string kernel SVM (*SVM-kmer*) (Arvey et al. 2012). Supplementary Table 3 lists 17 TFs for which ROCAUC was reported in (Arvey et al. 2012), where the mean accuracy across multiple cell lines was reported for each TF. We therefore compared the published accuracy with the mean *EMT* performance of the TF across only the cell types that were considered previously. As shown in Fig 2, in most cases, *EMT* outperforms *SVM-kmer.* DNAse hypersensitive (DHS) of a region represents its accessibility by DNA-binding proteins and previous studies have shown that integrating DHS with *in vitro* binding specificity can substantially enhance *in vivo* binding prediction (Arvey et al. 2012; Pique-Regi et al. 2011). Surprisingly, using pwm1k features 6 cases *EMT* outperforms even the model that integrated DHS with the kmer frequencies in the SVM (green). In a few cases (blue), SVM-kmer yields either comparable or improved predictability. Overall, the *EMT* models predict *in vivo* binding with a greater accuracy than a non-ensemble SVM approach represented by *SVM-kmer* (Arvey et al. 2012).

**Figure 2:**
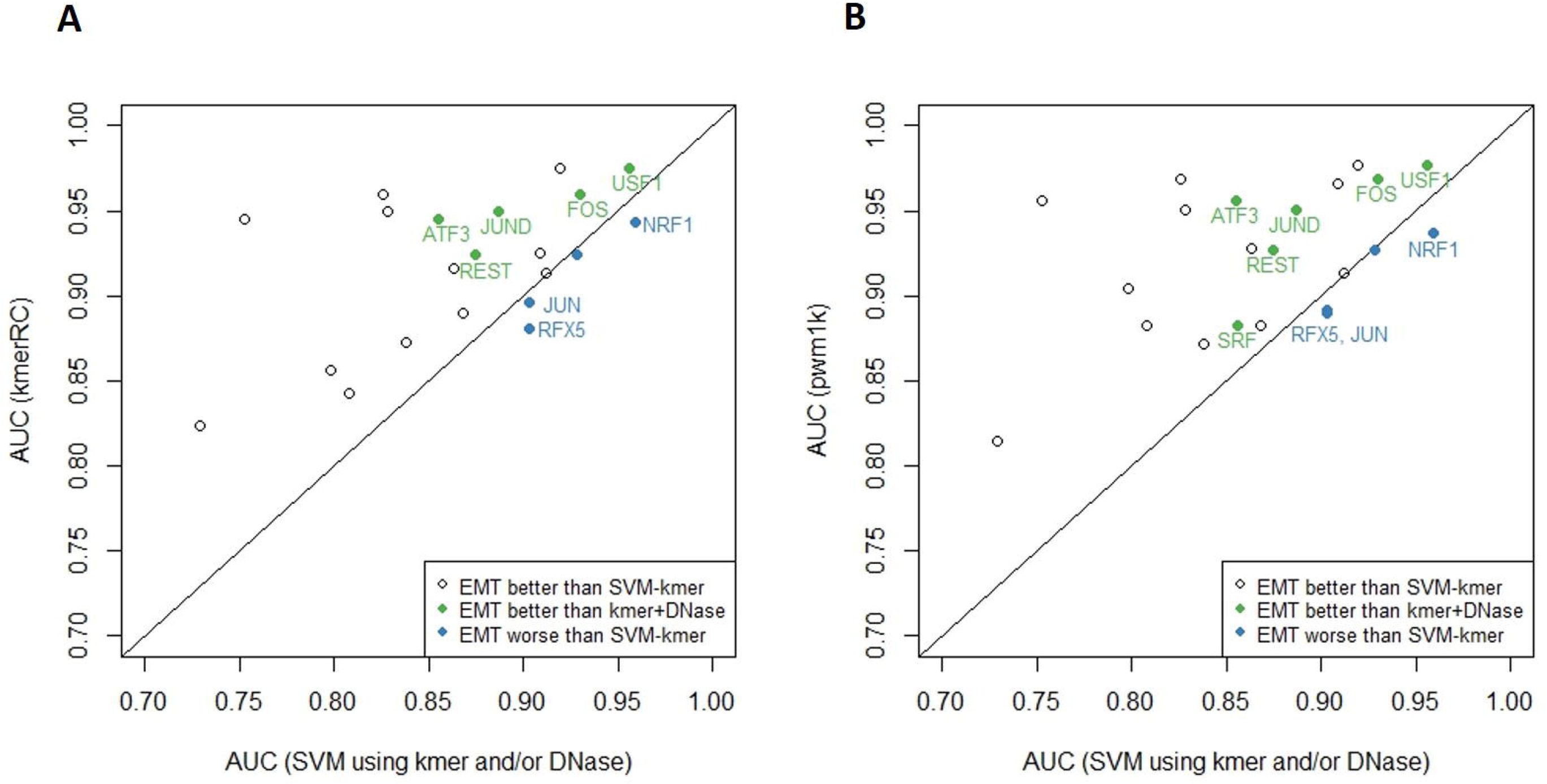
Prediction accuracy comparison of EMT against svm-kmer and svm trained using both kmer and DNase (kmer+DNase), where (A) EMT is trained using kmerRC features, and (B) EMT is trained using pwm hits with lkb stringency (*pwm1k*). Each point represent a TF. Except for 3~4 TFs (blue), EMT outperform svm in all other cases. For some TFs (green), sequence based EMT outperforms sequence+chromatin based model as well.

In sum, we have described a novel ensemble-based approach to in vivo binding modeling and established its superiority relative to SVM-kmer across a wide variety of TFs and cell types.

### TRISECT reveals intra-cell type heterogeneity and inter-cell type sharing of binding rules across cell types

The architectural difference and performance advantage of *EMT* relative to *SVM-kmer* suggests that *EMT* might be better able to exploit heterogeneous binding rules across the genome dictated by different combinations of interacting TFs. For each TF, we clustered the sub-models obtained from different cell types. As an illustrative example, Fig 3A-B show the cluster-membership matrix for TF *ATF3* for number of clusters *k* = 16 and 20. Fig S1 includes such mapping for all other TFs for *k* = 16. We found both cell type-specific (Fig 3B, cluster #6) and ubiquitous (Fig 3C, cluster #20) clusters. Examining the cluster mapping for all TFs (Fig S1), a wide range of patterns emerge: for certain TFs most clusters map to single cell type, suggesting cell type-specific binding modalities of these TFs (*EP300, JUN*), while certain other TFs have ubiquitously applicable binding rules, such as *YY1* and *TBP*, suggesting cell type independent binding rules and, presumably, function. Importantly, many clusters consist of sub-models from multiple, but not all, cell types. We ensured that inter-cell type sharing of *in vivo* binding rule is not simply due to shared binding loci across cell types (Supplementary Notes & Fig S2). Subsequent analyses are based on *k* = 16; reasons for this choice are discussed in Supplementary Notes & Fig S3).

**Figure 3:**
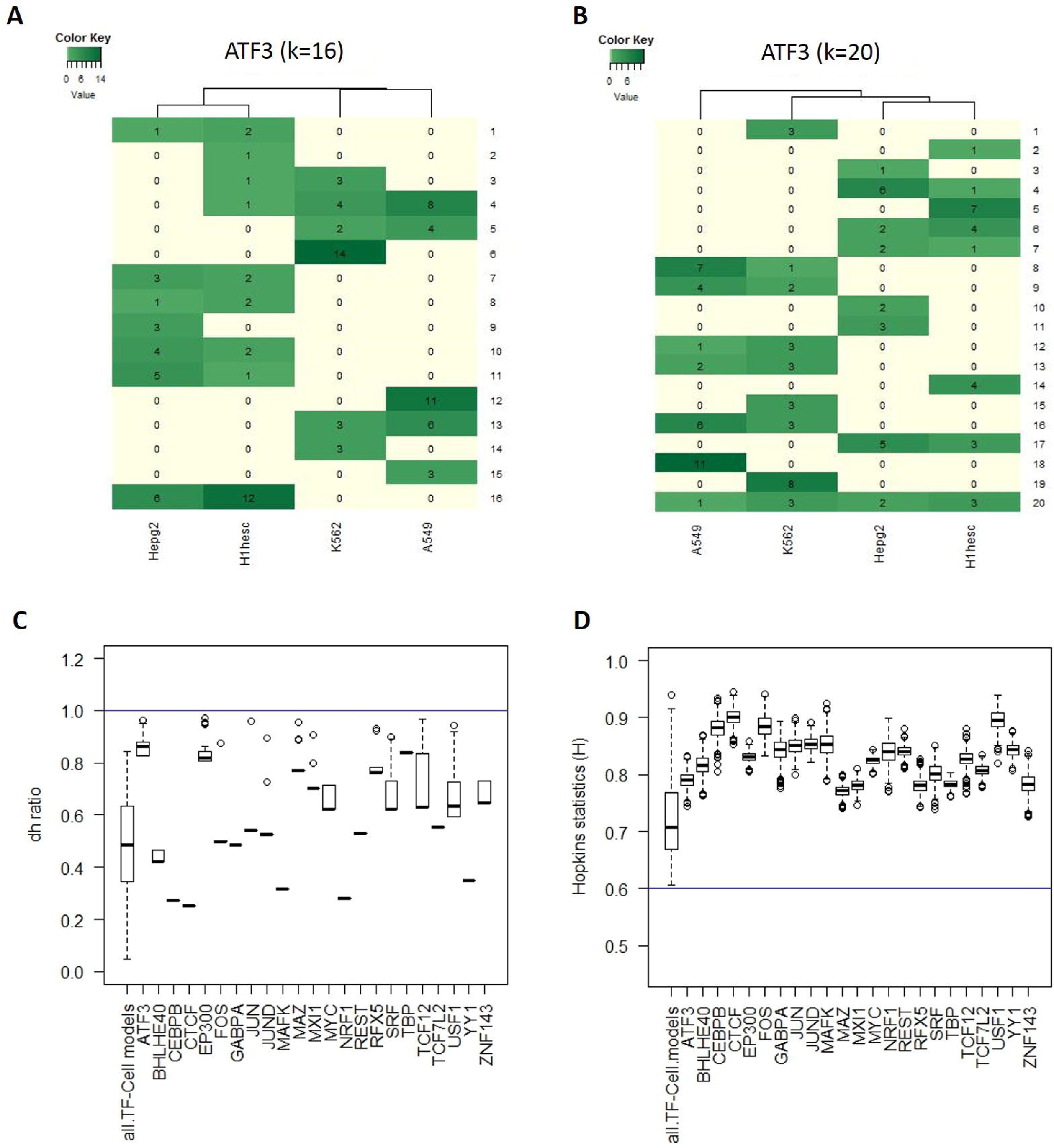
Assessing the existence of sub-models shared across cell types. (A&B). Cluster membership matrix using k-nearest neighbor clustering. Each row represents a cluster and column represents a cell type. Each element in the matrix denotes the number of sub-models in the cluster coming from each cell type. Some clusters consist of sub-models from multiple cells (cluster#20 in B), while some other consist of sub-models from a single cell type (cluster#6 in A). (C&D). Boxplot of *dh-ratio* and *Hopkins statistic* for 135 TF-cell pairs based on sub-models of TF-cell type pair, and pooling all sub-models for each TF.

It is possible that *EMT* can falsely yield multiple sub-models, even in absence of heterogeneity, and those sub-models can be falsely clustered. We ascertained heterogeneity across sub-models for a TF from multiple cell types using a *Dudahart test* (Duda et al. 2001) and assessed the clustering tendency of the sub-models in the d-dimensional feature space using *Hopkins statistics* (Jain & Dubes 1988). The *Dudahart* test verifies whether or not a set of data points should be split into two clusters from the estimate of within-cluster sum of squares for all pairs of clusters versus overall sum of squares; the ratio of the two sum of squares is quantified as the *dh-ratio.* On the other hand, the *Hopkins statistic (H)* compares the nearest neighbor distribution for a random set of points to the same distribution for the clustered sub-models (M&M). A value close to 0.5 indicates the sub-models are random set of points with no clustering, a value close to 1 indicates that they form a cluster. Fig 3C-D summarize the *dh-ratio* and *Hopkins statistic* respectively for 135 TF-cell pairs based on sub-models of TF-cell type pair, and for each TF after gathering all sub-models under a TF. We found that in all cases the *dh-ratio* is lower than 1 rejecting homogeneity (Fig 3C) and the set of sub-models form clusters (Fig 3D). All tests done for the analysis are significant (p-value <0.001) (M&M). Together, the *Dudahart test* and *Hopkins statistic* strongly suggest that the sub-models are distinct and clusterable, i.e., TF binding rules are heterogeneous and partly shared across cell types.

Next we assessed the functional underpinning of shared binding rules across cell types. Specifically, we assessed whether two co-clustered loci from different cell types (i.e., obeying similar binding rule) are functionally associated relative to loci from the same cell type but belonging to different clusters, i.e., obeying different binding rules. We devised a cluster-specific scoring of each binding sequence and assigned each binding site in each cell type to one or more clusters (M&M). As per convention, we assigned each binding site to the nearest gene as a potential transcriptional target; 95% of the target genes were within 100 kb from the binding site (median distance 4.5 kbp) (Fig S4). To assess functional coherence of a cluster, we determined the fraction of gene-pairs in the cluster (regardless of cell type) that participate in the same pathway as compared to all pairs of target genes within each cell type, and assessed the significance of enrichment using Fisher test. Likewise, we also estimated the expression coherence of genes within a cluster (M&M). As shown in Fig 4 and S5: ~40% (respectively, “‘18%) multi-cell type clusters show significantly higher (p-value < = 0.05) expression-coherence (respectively, pathway-coherence) than the background (expectation is 5%). Moreover, the pathway and expression coherence are highly correlated across clusters (spearman correlation=.56, p-value=0.02). We conducted the same set of tests for random clusters of same size as real clusters. In both cases, the coherence was no greater than the null expectation (Fig 4A-B). In Supplementary Tables 5a-b, we catalogue all the clusters with mapped target genes and their enriched GO terms.

**Figure 4:**
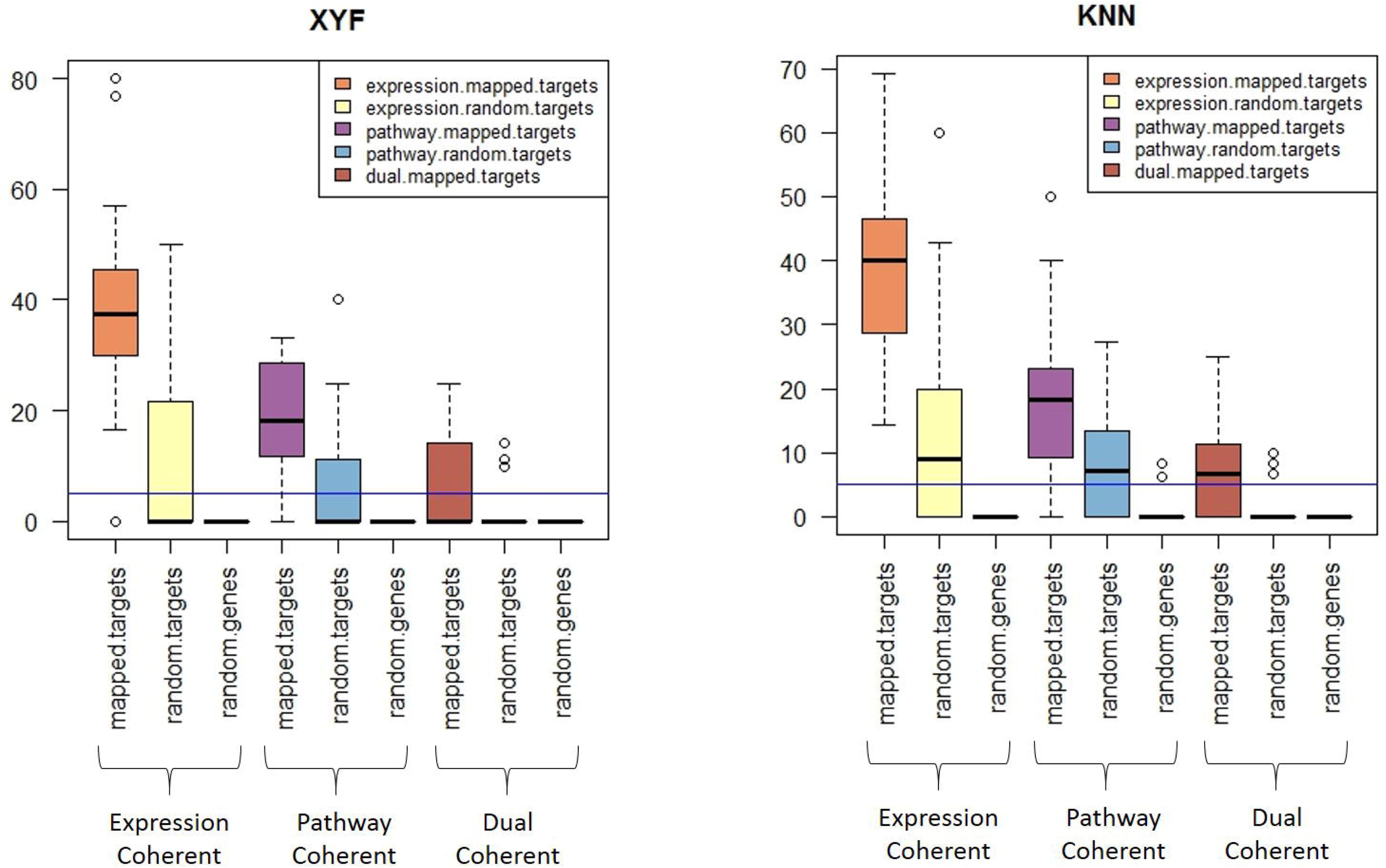
Functional and Expression coherence of sub-model clusters. (A&B) Fraction of multi-cell clusters found to be coherent using k-nearest neighbor (KNN) and XY-Fused (XYF) self-organizing map respectively. Mapped.targets denotes when genes are assigned to cluster based on *TRISECT* pipeline, random.targets indicates the clusters consisting of random genes among all targets and random.genes indicates the cluster consisting of random genes.

Taken together, these analyses support existence of heterogeneous sets of rule governing *in vivo* TF binding and that subset of rules are shared across cell types with functional implication.

### The role of interaction partners in a TF’s binding occupancy across cell types

By using 981 PWMs for a comprehensive set of vertebrate TFs as the basis for features, *EMT* implicitly incorporates the contributions of interaction partners in predicting *in vivo* binding of the reference TF. To quantify the contribution of interacting motifs, we repeated the *EMT* training and testing using only the PWMs corresponding to the reference TF. Individual TFs have multiple motifs reported in the literature (ranging from 1 to 8, with a median of 3; Supplementary Table 6), which can differ substantially from each other with potential functional implications (Bulyk et al. 2002; Hannenhalli 2008); we refer to these motifs as the *reference motifs*, and the *EMT* model utilizing only the reference motifs as the *NonInteraction* model and to contrast we refer pwm1k model as *Interaction* model. Supplementary Table 7 shows the prediction accuracies for the *Interaction* and the *NonInteraction models;* the diagonal elements represent the cross-validation accuracies within a cell type, while the offdiagonal elements represent the accuracy when *EMT* is trained on one cell type (row) and tested on another (column). Comparing the diagonal elements for the two models (summarized in Fig 5A), it is evident that *Interaction models* have higher predictive accuracy than *NonInteraction* models, which is consistent with the expectation that *in vivo* binding of a TF relies on interactions among several TFs.

**Figure 5:**
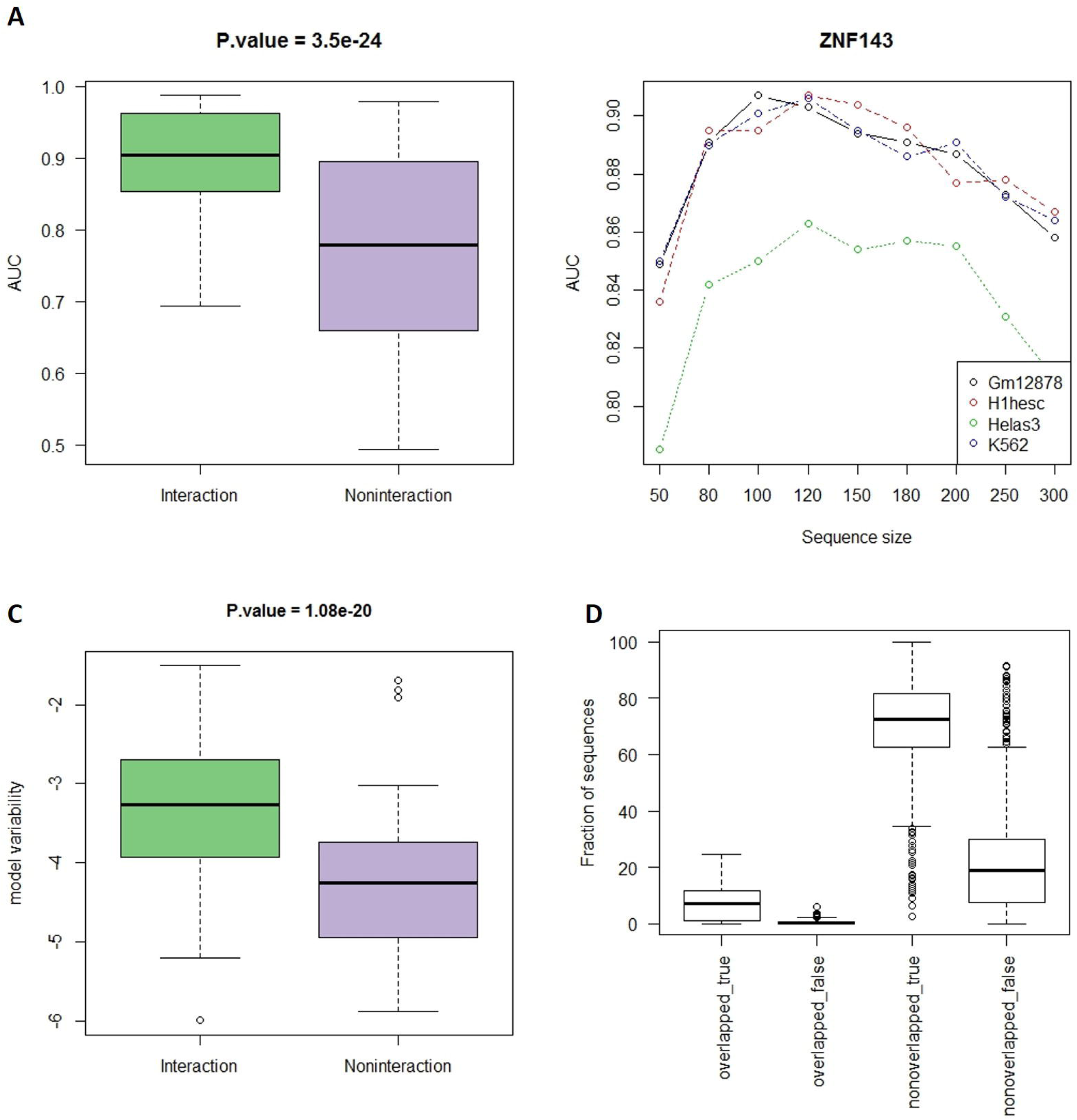
Association between number of interaction partners and model-accuracy. (A) The trend of model accuracy with increasing sequence size for TF ZNF143 (selected arbitarily for illustration). (B). Comparison of cross-validation prediction accuracy for *Interaction* and *NonInteraction* models. (C). Comparison of model variability in log scale (cross-cell type performance variability) for *Interaction* and *NonInteraction* models. (D). Distribution of the fraction of test sequences that fall in one of the four categories: Overlapped_true (respectively, overlapped_false) denotes the correctly (respectively, incorrectly) classified sequences having at least 50% overlap between the training sequences in one cell type and the test sequences in another cell type. Nonoverlapped_true (respectively, nonoverlapped_false) denotes correctly (respectively, incorrectly) classified sequences that do not overlap with any sequence in the training set.

Next, we conjectured that in the *Interaction* model, allowing for greater numbers of partners allows learning of more complex binding rules and increase binding prediction accuracy. We therefore assessed the effect of the length of the region flanking the binding site on prediction accuracy (M&M). We note that beyond 100bp, due to narrowing of the gap between the foreground and the background region, the discrimination accuracy is expected to decrease. Despite this, in some cases (Fig 5B & S6), the increase in ROCAUC beyond 100bp suggests that a larger context may be necessary in these cases to capture the binding rules. Nevertheless, we chose a sequence context of 100bp to make our model comparable to the previously published *SVM-kmer* (Arvey et al. 2012).

For a given TF, we also quantified the variability of the model accuracy in different cell types (M&M). We expect a model that relies on cell type-specific interaction partners to be more variable in its performance accuracy than the one that relies only on the reference motifs. This expectation is borne out in our analysis (Fig 5C). This suggests that part of the sequence information for *in vivo* binding is encoded by the TF’s own motifs and this does not vary substantially across cell types, while the additional context- and interaction-dependent part does. However, the small variability in cross-cell type prediction accuracy when using *NonInteraction* model is likely to come from the heterogeneity of binding motifs for a TF. We quantified the inter-motif divergence for each TF as either the number of motifs annotated for the TF, or motif-divergence defined over all motifs-pairs) (M&M). We found that the *NonInteraction* model performance variability is positively correlated with both measures of motif divergences (Spearman correlation=.63, 0.67; p-value=1.2e-3, 6.3e-4 respectively).

For the *Interaction* model, the off-diagonal elements in Supplementary Table 7 show relatively high cross-cell type performance accuracy, suggesting that the binding ‘rules’ are shared between cell types. We ensured that the high cross-cell type prediction accuracy is not simply due to shared sequence information, i.e., the genomic loci on which the model was trained in one cell type does not substantially overlap with the loci tested in another cell type. Overall, across all TFs and all pairs of cell types, the fractional overlap in genomic loci ranges from 0 to 10%, with a mean and median of ~4% (Fig 5D). This suggests that it is the binding rule, independent of specific sequence instances, that is shared across cell types.

Furthermore, we found that when using the *Interaction* models, the cross-cell type accuracy is symmetric (Spearman correlation of upper and lower triangle in Supplementary Table 7 is 0.68, p-value 9.5e-53). In other words, a high (respectively, low) accuracy in cell type *Y* using *EMT* trained on cell type *X* implies a respectively high (respectively, low) accuracy in cell type *X* using the model learnt from cell type *Y.* This further supports that the interaction-dependent (therefore genomic-context dependent) binding rules are shared across cell types. In stark contrast, there is a lack of symmetry in cross-cell prediction accuracy when *NonInteraction* model is used (Spearman correlation = 0.04, p-value 0.4).

In sum, our analyses suggest that the cell type-specific TF interactions play critical role in determining cell type-specific *in vivo* binding. In addition to that, these revealed by *EMT* might be responsible for cell specific binding of the reference motifs.

### TRISECT reveals putative co-factors providing insights into cell-specific biological roles of a TF

Our results so far suggest that cell type-specific co-factors of a TF are a major driver of variability in the *in vivo* binding rules across cell types. To further probe into the functional implications of cell type-specific co-factors, for each reference TF, we identified its cell type-specific co-factors using the feature importance of the corresponding motif as estimated by the model. To minimize redundancy, we excluded motifs with substantially high co-occurrence frequency with at least one of the reference motifs (M&M). To further minimize false positives, we assessed the enrichment of motif occurrence near the cell-specific ChIP-Seq peaks of the reference TF relative to background and retained only those putative co-factor motifs that were significantly enriched (odds ratio > 1.2 and p-value < 0.05, M&M). The choice of enrichment odds ratio threshold is rationalized in Fig S7, which shows that increasing the threshold would result in a loss of information for some TFs e.g. REST.

Several lines of evidence support the cell type-specific co-factors for a TF identified by *TRISECT.* First, we found that for ~70% of the models, the putative co-factors are enriched for either heterodimerizing TFs or for the TF family that the reference TF belongs to (M&M & Supplementary Table 8). The enrichment of same family as that of reference TF is consistent with the fact that TFs forms dimer with other TFs preferably from same family (Amoutzias et al. 2008; Dror et al. 2015). We also performed protein domain enrichment analysis (Supplementary Table 9) using DAVID tool (Huang, Brad T. Sherman, et al. 2009; Huang, Brad T. Sherman, et al. 2009), and found that more than 80% of enriched domains are involved in homo- or hetero-dimerization consistent with Supplementary Table 8.

Second, we expect putative co-factors to be expressed at higher level in the specific cell types where they are deemed as co-factors. For each co-factor (excluding ubiquitous co-factors), we determined the log-fold difference in expression between the cell types where it is identified as co-factor relative to cell types where it is not (M&M). The distribution of log fold changes of the co-factors are compared with a control set of fold ratios as presented in Fig 6A. For most TFs, the co-factors show significantly higher expression in the relevant cells. This is not true only in 5 cases. Among these, *CTCF* is known as cell type-independent TF and for two of them (*GABPA* and *NRF1*) we show below, via an independence test, that they show higher cell independence than other TFs.

**Figure 6:**
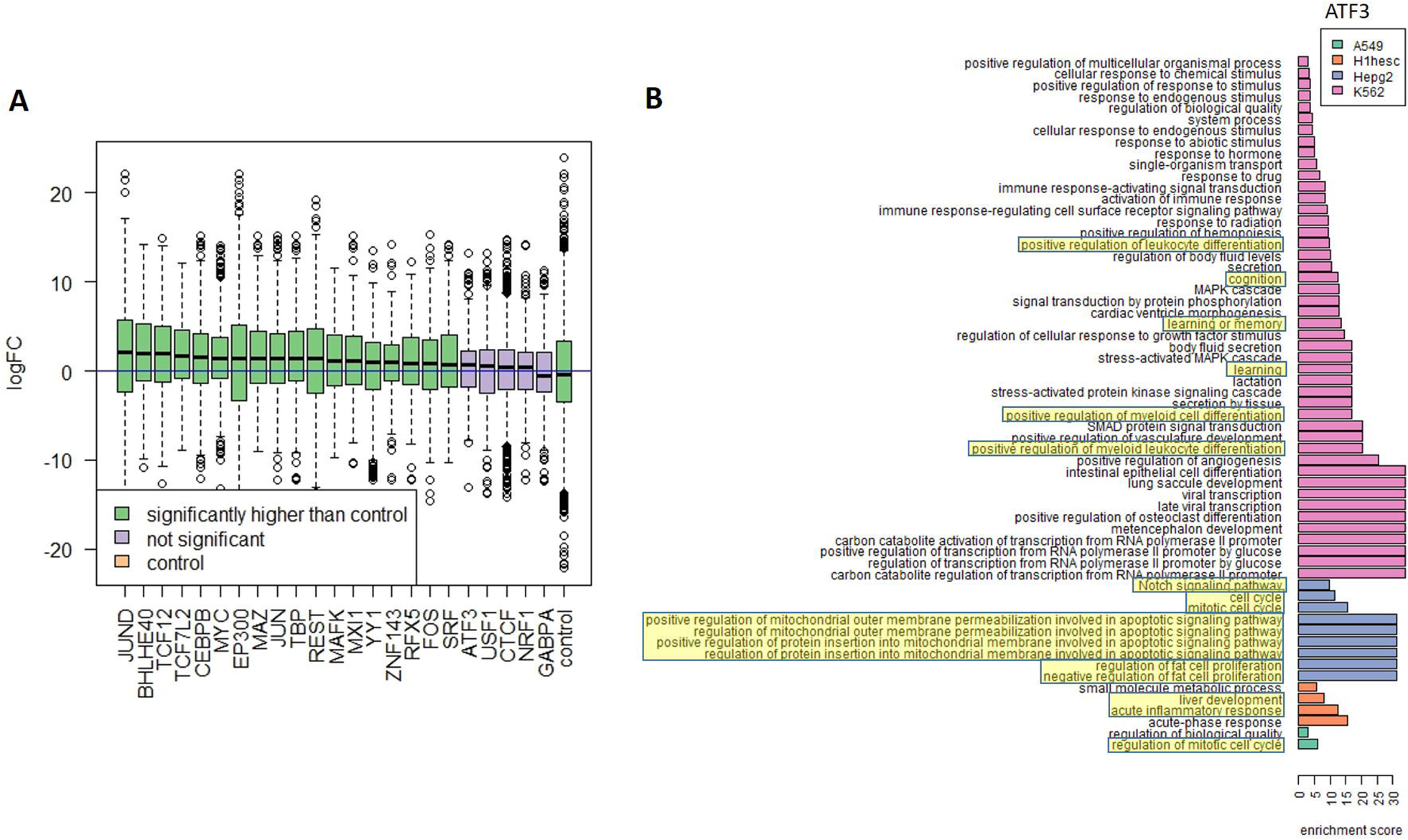
Functional validation of putative co-factors. (A). Identified co-factors have higher expression in the cell lines they are detected in. For a TF motif detected as a co-factor in *n* cell lines, and not in another *m* cell lines, we calculated fold difference in the TF’s expression between the two sets of cell lines. Each boxplot corresponds to all co-factors of a TF in X-axis. (B). As an example, for ATF3, GO enrichment analysis of co-factors in four cell types recapitulate the known cell type-specific biological roles.

Third, for each TF’s cell type-specific co-factors, we performed biological processes GO term enrichment analysis using the Gorilla tool (Eden et al. 2009) relative to all 981 motifs as the background. We found significant differences in function among co-factors for a TF in different cell types. Remarkably, the biological processes can vary across cell types while still being functionally related to the reference TF. As an illustrative example, Fig 6B shows the enriched BP (false discovery rate < = 10%) for ATF3 in 4 cell types. ATF3 is a stress-inducible TF involved in homeostasis (Allen-Jennings et al. 2001; Tanaka et al. 2011), specifically regulating cell-cycle, apoptosis, cell adhesion and signaling (Tanaka et al. 2011). We found that ATF3 co-factors are enriched for functions related to cell cycle and proliferation in 3 out of 4 cell lines. In stem cell, the identified co-factors are involved in liver regeneration and inflammatory response, consistent with previous studies showing direct link between ATF3 induction and liver injury and regeneration in mice (Chen et al. 1996; Su et al. 2002). Furthermore, enrichment of NOTCH and apoptotic signaling among co-factors in Hepg2 cell line is consistent with role of ATF3 in glucose homeostasis and other primary functions of the liver (Allen-Jennings et al. 2001). Surprisingly, we find enrichment of cognition, learning and memory among the co-factors in leukemia cell line. Since leukemia is a cancerous cell line, non-native gene expression is not unexpected (Lotem et al. 2004; Lotem et al. 2005). However, even though ATF3 is not known to play a direct role in neuronal function, a closely functionally and structurally related protein CREB has well documented role in neuronal activity and long-term memory formation in brain (Mayr & Montminy 2001), raising the possibility that either ATF3 has a hitherto unknown role in cognition or, alternatively, the same set of co-factors are involved in memory formation in conjunction with other TFs.

For other TFs, the enriched GO-terms at false discovery rate cutoff of 10% (enrichment scores ranges from 1.22 to 93.75 with a median of 7.44) are listed in Supplementary Table 10 with corresponding discussion based on literature survey is provided as Supplementary Notes. This can serve as a resource for further investigation into cell type-specific binding and function of a broad array of TFs. In Supplementary Tables 5a-b, we catalogue all the clusters with their specific TF interactions (M&M), and their enriched GO terms.

We noted substantial variability in the number of detected co-factors across cell types for a TF. Interestingly, a literature survey suggests that the cell types where the reference TF has specific function, the number of co-factors in that cell type is comparatively higher. For example, REST has well-known neuronal functions and its binding sites in neurons exhibit lack of cognate RE1 motifs (Rockowitz et al. 2014), suggestive of dependence on co-factors. Consistently, Sknsh (brain cancer cell line) has highest co-factor cardinality for REST. Similarly, JUN plays specific role in hematopoetic differentiation and we found that Gm12878 (normal blood cell line) has the largest number of co-factors (Liebermann et al. 1998). We reasoned that TF with greater cell type-specific roles would exhibit greater variability in co-factor cardinality. For each TF we measured the variability of its co-factor cardinality across cell types. As shown in Fig 7A, interestingly, TFs with ubiquitous and invariant roles such as TBP and CTCF have the least variable co-factor cardinality.

**Figure 7:**
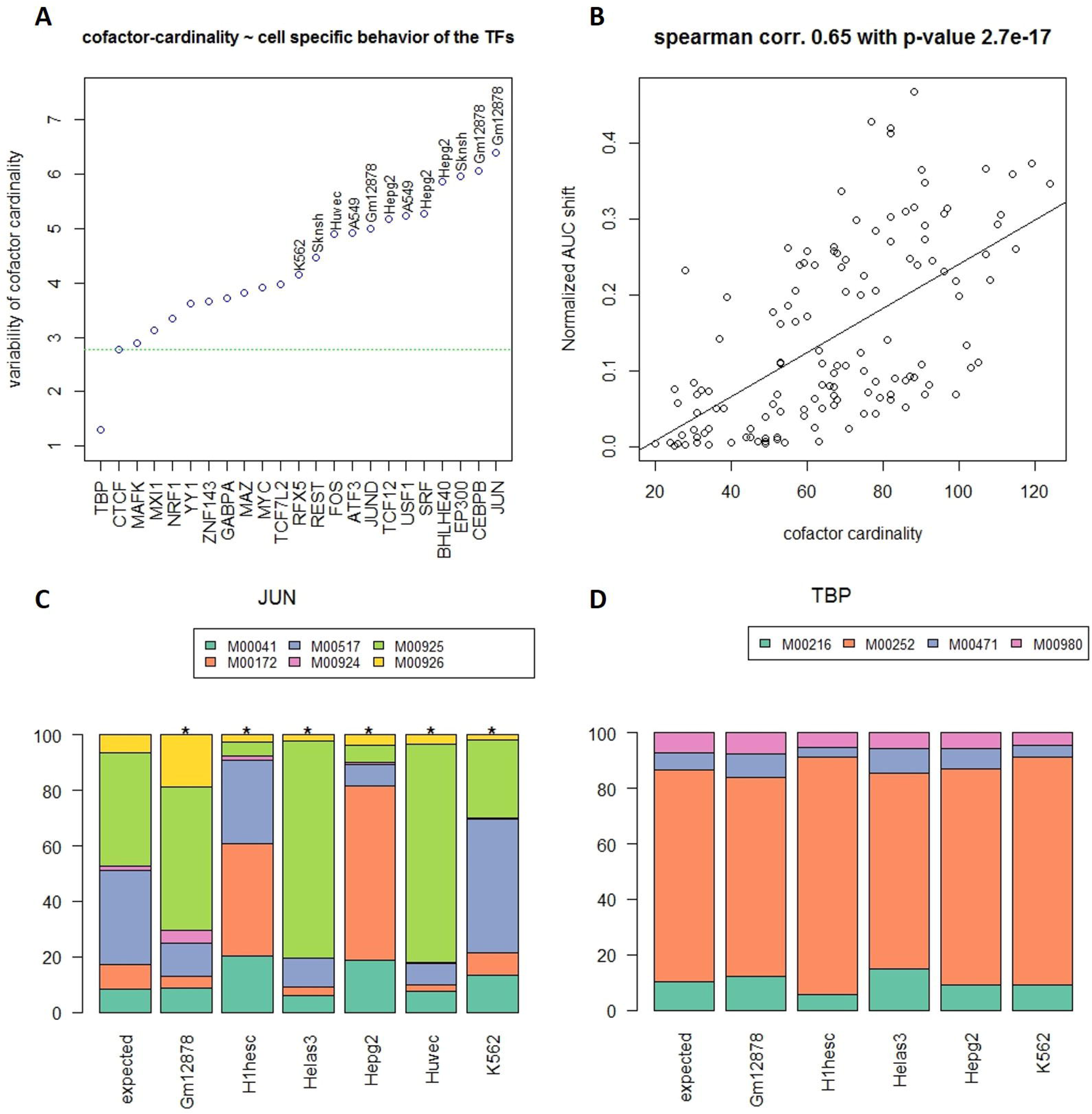
EMT model heterogeneity is associated with cell type-specificity of co-factors. (A) The plot shows for each TF the variability of co-factor cardinality across cell types. Each point is further labeled with cell type where the relevant TF has specific usage, based on literature and has largest number of co-factors. TBP and CTCF are the most ubiquitous TFs. (B) Normalized ROCAUC difference of *Interaction* and *NonInteraction* models for a specific TF-cell type pair correlates with co-factor cardinality. (C-D) Cross-cell type variability in motif usage for the reference TF in the *NonInteraction* model, for JUN and TBP as two extreme examples. JUN shows different binding specificity in different cell types, while TBP does not.

We also assessed whether the difference in prediction accuracy achieved by *Interaction* model and the *NonInteraction* model for a particular TF-cell type pair may reflect the TF’s dependence on co-factors. We measured the normalized distance between the performance (*performance distance*) of *Interaction* and *NonInteraction* model (M&M) and compared it with co-factor cardinality. As shown in Fig 7C, we found that the *performance distance* is positively correlated with co-factor cardinality (Spearman correlation = 0.65, p-value = 2.7E-17).

Previous studies have found that the DNA sequence specificity of a TF can be influenced by interaction with co-factors (Siggers et al. 2011; Slattery et al. 2011). Interestingly, a close inspection of the feature importance estimated by the *NonInteraction EMT* model shows that in different cell types different compositions of the reference motifs are utilized. Fig S8 presents all cell type-specific usage of a TF’s motifs; the cells where the motif usage is significantly different from expected usage are marked with asterisk (M&M). Notably, such diverse usage is observed using *NonInteraction* models, suggesting cell type-specific motif preference even without any modulation by the co-factors.

Taken together, the cell type-specific co-factors revealed by TRISECT are consistent with their cell type-specific expression and function and may be critical in modulating a TF’s cell type-specific biological function.

## Discussion

In this study, we have presented a novel ensemble-based framework –*TRISECT*, to investigate intra-cell type heterogeneity of *in vivo* TF binding rules and inter-cell type commonality thereof. To the best of our knowledge, this is the first study to show, based on a comprehensive analysis, that *in vivo* binding specificity rule is composed of multiple components, or sub-models, many of which are shared across multiple cell types. Tellingly, non-orthologous targets of binding sites across cell types governed by a shared binding sub-model exhibit a greater functional and expression coherence than targets of binding sites in the same cell type that are governed by different binding rules. For each TF, *TRISECT* identified cell type-specific co-factors that are supported by gene expression data and literature studies supporting their cell type-specific function. As a useful functional resource, for 23 TFs included in this study, we provide a catalogue of clusters of shared sub-models, along with their putative cell type-specific targets, the co-factors characterizing the cluster and their function.

Our ensemble model not only outperformed the previously reported sequence-based discriminative model (*SVM-kmer*), but in several cases it outperformed the model that utilizes the chromatin accessibility in addition to the sequence flanking the binding site (Arvey et al. 2012); paradoxically, some of the TFs (e.g., JUND) whose *in vivo* binding were deemed to depend less on the sequence context and more on the chromatin accessibility by the previous SVM approach were found to be adequately modeled by sequence alone when using the EMT approach. Taken together with our observation that these TFs depend on a large number of cell-type exclusive co-factors for their *in vivo* binding, these results suggest that cell type-specific chromatin accessibility is captured, to some extent, by binding sites for cell type-specific co-factors, shown independently by recent work (Whitaker et al. 2015; Benveniste et al. 2014). Apart from the modeling approach of a TF’s *in vivo* binding specificity, our study differs from Arvey et al (Arvey et al. 2012) in several other aspects. In discussing cell type-specificity, the previous study compared the models only in two cell types – GM12878 and K562, while we have investigated in-depth the cell type-specificity of *TRISECT* across 4-12 cell types. While the previous work primarily discusses cell type-specificity and ubiquity of their models, by clustering the cell type-specific sub-models, our work investigates the extent of shared binding rules; cell type-specificity and ubiquity are extreme cases thereof. In addition to cell type-specific variability in proximal co-factors, we investigated in much greater depth than the previous work the cross-cell type variability in the preferred motif for the reference TF. Together, these novel aspects of our study adds to the knowledge of sequence information that specify a TF’s *in vivo* binding in various cell types.

Another recent study (Dror et al. 2015) aiming to decipher the determinants of *in vivo* occupancy of a TF showed that TF binding specificity is influenced by nearby homotypic sites (for the reference TF), the local nucleotide composition, and certain DNA physical properties. Moreover, a preferred *in vivo* binding in a homotypic cluster was shown to be related to a preferred nucleotide composition (GC-rich for zinc finger TFs and AT-rich for homeodomain reference TFs) in the flanking region of the binding site. These previous findings are consistent with the fact that the co-factors identified by *TRISECT* are enriched for same family of TFs as the reference TF and thus have similar preference for nucleotide composition as the reference TF. In the previous work (Dror et al. 2015), the accuracy in discriminating bound vs. unbound sequences after controlling for the presence of a putative site for the reference TF was modest (ROCAUC ~ 0.6). Whereas, we have shown that the motifs for the reference TF alone can discriminate bound from the unbound control sites with ROCAUC ~ 0.78, suggesting that the reference TF are most informative in determining in vivo binding, as also observed in Pique-Regi et al (Pique-Regi et al. 2011), and the additional power of discriminations comes from the presence of co-factor motifs, as suggested before (Arvey et al. 2012; Hannenhalli & Levy 2002), or from nucleotide composition and various DNA physical properties (Dror et al. 2015). Interestingly, DNA flexibility measured by propeller twist (el Hassan & Calladine 1996) is highly dependent on GC-content (Hancock et al. 2013), which in turn is related to motif composition, as we have noted. Overall, these seemingly independent properties (nucleotide composition and DNA physical properties on one hand and motif composition on the other) may be related. Specific advantage of an ensemble model based on motif composition is that apart from being highly accurate, it is functionally interpretable and provides insights into a TF’s cell type-specific functions.

Context-dependent function of a *cis* regulatory region requires binding of a specific combination of TFs. This modularity contributes to morphological evolution through changes in cis elements controlling transcription, while avoiding the pleiotropic effects of TF gene’s expression change (Prud’homme et al. 2007). Shared sub-models of TF binding rules across cell types, as revealed by *TRISECT*, may suggest shared history of cell types.

The ability of a TF to bind to diverse reference motifs and in conjunction, interact with diverse combinations of co-factors serves to enhance its functional repertoire across contexts (Meijsing et al. 2009; Arvey et al. 2012). Our analyses indeed reveal cell type-specific preference for the reference motif as well as the cell type-specific interaction partners of a TF. We found that the expression of cell type-specific interaction partners to be higher in the cell types where they are expected to interact with the TF and their function are consistent with the context based on the literature. Thus our study provides further support for a TF’s cell type-specific functions, and more importantly, enables further investigation into the mechanisms underlying a TF’s diverse cell-specific functions.

## Methods

### Data Processing

We downloaded the ChIP-Seq peaks or 23 TFs from ENCODE (Supplementary Table 1). For each TF we selected only those cell lines for which narrow-peak data was available. We chose the more stringent of the two criteria - top 5000 most significant peaks, or FDR q-values<0.2 to select binding sites (Arvey et al. 2012). Relative to the center of ChIP-Seq peaks, the DNA regions of length 100bp were identified as the foreground. As negative control, we sampled flanking regions of 100bp from 200bp away from the positive sequences. Moreover, control sequences overlapping with any peak were excluded. Due to the proximity of the negative examples, both foreground and background are expected to have similar GC-composition (Arvey et al. 2012) and chromatin accessibility. However, we explicitly controlled for the GC composition using sequence set balancing technique when comparing the foreground and the background (Whitaker et al. 2015). We discarded any cell line resulting in fewer than 4000 sites.

### Learning EMT

We considered three types of feature set for the sequence specificity model: (a) kmers - frequencies of 4096 6-mers in the 100bp sequence, (b) kmerRC - frequencies of 2080 6-kmer groups equating a k-mer and its reverse complement, and (c) pwm***lk***–we take all the positional weight matrices (pwm) from TRANSFAC 2011 as the features and get the motif hits using PWMSCAN (Levy & Hannenhalli 2002). The feature value is the sum of pwm-score (-log10(hit score)) obtained from the PWMSCAN; we took the log of feature values to compensate for the skewed distribution of the number of binding sites. Here, ***lk*** refers to the PWM hit threshold (hit expected every *l* kb on average in the genome); we used *l* = 1/2/5/10kb.

We chose Adaptive boosting (Freidman 2008; Friedman 2002) as our composite model where each sub-model within the ensemble is a decision tree and each decision tree is constructed based on a bootstrap sample. We used the Adaboost framework implemented in R gbm package (Ridgeway 2015). In the framework, Huber loss function is selected to reduce over-fitting. We estimated the classification accuracy of the model based on 25% held out data set, while 75% data were being used to build each tissue-specific model.

### Model conversion, Dudahart test and Hopkins statistics

Each sub-model is represented by a point in a *d*-dimensional space. Each dimension denotes a feature and the value along the dimension indicates the importance of the feature for the sub-model. Therefore, each model (consisting of multiple sub-models) can be represented as a set of points in an *n*-dimensional space where *n* ≤ 981. For a model, the feature importance was measured based on the prediction performance improvement by evaluating predictions on an out-of-bag samples. We modified the gbm package (Ridgeway 2015) implementation of feature-importance to accommodate the calculation for single tree or the sub-model in question. In other words, we determined the contribution of a single tree (sub-model) in prediction performance improvement using the same out-of-bag samples. We disregard the features which do not contribute to any sub-model. We measured dh-ratio (ratio of within-cluster sum of clusters and overall sum of squares) for all cluster pairs, based on either cell type-specific set of sub-models, or the pooled set of sub-models across all cell types for a TF. While calculating dh-ratio, K-nearest neighborhood (KNN) approach was used for clustering. Since the final output of KNN depends on initial random set of centers, the dh-ratio calculation was repeated 1000 times to ascertain robustness. We noted that all test results were significant (p-value < 0.01).

To measure Hopkins statistics (H) the sub-models are again represented as a set of points. H is defined by the following.

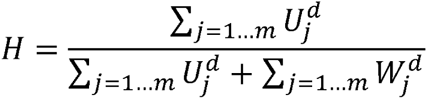

*W_j_* are the nearest-neighbor distances of *m* randomly chosen points (sub-models), which demarcate the sampling window. *U_j_* are the minimum distances of the sub-models from *m* random points in the sampling window. To define the sampling window, we either took 25 to 75 percentile of the feature values or from δ to max.value-δ along each dimension, where δ denotes the standard deviation of the feature value (Dubes & Zeng 1987; Zeng & Richard C Dubes 1985; Zeng & Richard C. Dubes 1985). To estimate p-value, we repeat the above procedure 1000 times and measured the H value. The p-values ranges from 0.026 to less than 0.001.

### Clustering sub-models

For a TF, we obtained sub-models in all cell types, and then clustered all sub-models using K-nearest neighbor (KNN), where each sub-model is an instance and the features of the instances are individual feature-importance obtained in the context of respective tissue-specific model. Before feeding into the KNN, we remove all the features whose cumulative importance over all sub-models is zero. The sub-models are also clustered using XY-fused version of self-organizing map (Melssen et al. 2006) from kohonen R package (Wehrens 2015). To make it comparable to KNN, we assumed 100% weight for X map, i.e. sub-models will be clustered without preexisting label of which sub-models belonged to which cell.

### Assignment of sequences and target genes to the clusters

A cluster of sub-models can be viewed as a new ensemble. We scored each binding site sequence against each cluster, and a sequence is assigned to a cluster when it is scored above a threshold (of 1) by the cluster. The choice of the threshold was based on the rationale that the *intercept* (Ridgeway 2015) of tissue-specific models are ~1, and for a high-confidence positive sequence, the model-score should be greater than the intercept. Each bound sequence (from all cell lines) is mapped to a set of clusters. For each bound sequence, the nearest gene on the genome is considered to be its putative target, as per convention (Zhu et al. 2010). Hence, each cluster corresponds to a set of target genes coming from different tissues. We arranged the target genes into an *M*-by-*N* array, where *M* is the number of cell lines and *N* is the number of clusters. The enriched pathway among the target genes of each cluster was determined using clusterProfiler R package (Yu et al. 2012).

### Measuring functional and expression coherence using Fisher test

We downloaded the KEGG pathways (www.genome.jp/kegg). We use the following contingency table to determine whether the target genes from different cell lines that are assigned to the same cluster are more functionally related than the target genes coming from the same tissue but from different clusters.

| Gene pair across |  | Cluster<br>(Foreground) | Cell line<br>(Background) |
| --- | --- | --- | --- |
| In same<br>Pathway? | Yes | $a$ | $c$ |
| | No | $b$ | $d$ |

In the *M*-by-*N* target gene array, we compared all gene-pairs along columns from different rows (same cluster, different tissues) and the gene-pairs along rows from different columns (same tissue, different cluster) as the background. Then we apply the Fisher exact test in a cluster-centric fashion by comparing the fraction of foreground gene-pairs in the same pathway relative to the background.

Expression coherence tests were designed similarly, based on the following contingency table.

| Gene pair across |  | Cluster<br>(Foreground) | Tissue<br>(Background) |
| --- | --- | --- | --- |
| Co-expressed? | Yes | *a* | *c* |
|  | No | *b* | *d* |

A gene-pair is considered co-expressed if both of the genes are turned on (RNA-seq log2CPM > 1) in their respective tissues; CPM stands for Counts per Million. CPM, instead of the standard FPKM measure to quantify gene expression suffices for our purpose as we only compare a gene’s expression across samples, and not with other genes in the same sample. We showed similar trend of expression coherence with different expression threshold (log2CPM>=5) (Fig S5).

### Model variability, and Motif-divergenece

Model variability is defined by its normalized-predictability across cell lines. For each model, n ROCAUC values are obtained on held-out dataset of n cell-lines. Cross-ROCAUC values are normalized by self-ROCAUC value. Mathematically, 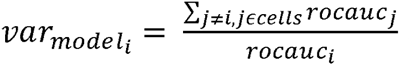.

Motif-divergence is defined by the following equation, *motif.* 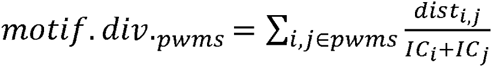. Here, *dist_i,j_* = 1*/similarity_i,j_* and *IC_i_* is the information content of ith motif. Similarity between two pwms is calculated following the normalized version of the sum of column correlations (Pietrokovski 1996).

### Identification of co-factors

EMT provides importance of all features in discriminating the foreground from the background. We retained all features with nonzero importance. From the initial set, we removed any motif that has 60% pwm-similarity (consensus overlap) for at least 50% of the binding site locations with any of the reference motifs. Next, we calculated enrichment of the motif in the foreground binding sites relative to control sites. We retained the motifs with greater than 1.2-fold enrichment and p-value < = 0.05. The resulting motifs were considered as cofactor. For further analysis, we considered tissue specific cofactors by removing common motifs across tissue. For *unique-relaxed* set we excluded co-factors that are common across all cell-lines, and for *unique-strict* set co-factors common to any two cell lines were excluded. The functional tissue-specificity measure for a TF is determined using the cardinality-variability of unique-strict co-factors.

### Gene expression and differential gene expression

For gene expression, we used RNA-seq data downloaded from ENCODE (Supplementary Table 4). For each tissue, we obtained between 2 and 4 RNA-seq samples depending on the availability and obtained the number of reads aligned to the gene. We corrected for batch effect using ComBat tool (Leek & Storey 2007). To estimate differential expression between two set of cell lines (those in which a TF is deemed a co-factor, and those where it is not), we used linear model from R package, limma (Smyth 2005).

### Enrichment of same family TFs and heterodimerizing TFs

We collected the family name of each PWM and the list of heterodimerizing PWMs based on semi-automated inspection of TRANSFAC 2011 annotations, based on keywords and further reading of the description. For hyper-geometric test of family-enrichment, we compared how many co-factors belong to the family of reference motifs relative to the 981 motifs. Heterodimer enrichment was tested similarly.

### Cluster specific TF-interactions mapping

Cluster-specific co-factors are identified by treating a cluster as a new ensemble of sub-models. We computed an aggregated relative importance of the features, considering the decision trees of the new ensemble corresponding to a cluster. Since the set of decision tree has been changed from the original set of trees from the EMT, some of the detected co-factors may be false positives. We took the intersection of the features (with non-zero importance) with the ‘enriched-nonoverlapped’ (or ‘distinct-relaxed’ or ‘distinct-strict’) co-factors of the original EMT. The corresponding enriched GO terms are determined using a R package called clusterProfiler (Yu et al. 2012).

### Tissue-specific pwm for the reference TF

We obtained relative feature importance of the reference motifs from the *NonInteraction* models and compared them with random expectation. To calculate the random expectation, 1000 *NonInteraction* models are learned based on randomly sampled 4k sites from among all binding sites across cell-lines. From 1000 models 1000 relative feature importance is calculated. Each set of relative importance is assumed a point in p-dimensional space where p is the number of reference motifs. We considered the relative importance vectors as data points from multivariate normal distribution and for each vector we calculated the Mahalanobis distances from the centroid which follows a chi-square distribution (Slotani 1964). The degrees of freedom (d) for the chi-squared distribution is determined using maximum likelihood estimate and a P-value is generated from a chi-square distribution function of d degrees of freedom.

## Supplemental information

Supplemental Figures, S1-S8

Supplementary Tables, 1-10

Supplementary Notes, 1-3

## Authors’ contributions

S.H. conceived the project. S.H. and M.S. designed the analyses in consultation with H.C.B. M.S. performed the analyses. S.H. and M.S. wrote the manuscript with help from H.C.B.

## Acknowledgements

This work was supported by NIH R01GM100335 to S.H. and NIH R01HG005220 to H.C.B. We thank Justin Malin and Avinash Das for helpful comments and suggestions. M.S. wishes to thank Justin Malin and Hiren Karathia for extensive discussion on heterodimerization and biological processes.

